# Neglecting movement-disease feedback biases wildlife epidemic forecasts

**DOI:** 10.64898/2026.07.30.741907

**Authors:** Yun Tao, Rodrigo L. Campos, Mark Q. Wilber, Liliana C. M. Salvador

## Abstract

In wildlife populations, movement drives host contact networks and disease spread. Growing evidence from veterinary science and experimental biology shows that infection commonly modifies host movement itself, a widespread feature of host biology. Yet conventional wildlife disease models treat movement as independent of infection, a mechanistic omission that could bias epidemic forecasts and management decisions. Here, we introduce a general framework in which recent infection temporarily alters a host’s home-ranging behavior and where pathogens are shed, integrating movement and disease ecology through spatially explicit, individual-based simulations. We show that infection-modified movement (IMM) responses, including anorexia, disorientation, and hypokinesia, can enhance or suppress epidemics, and most responses reverse direction depending on environmental conditions, grouping structure, and social behavior. We further show that models without IMM alternately underestimate and overestimate epidemic size as ecological context changes. Because our model parameters map onto quantities routinely estimated from tracking and surveillance data, our framework enables movement-disease feedback to be measured and incorporated into wildlife disease models.

## Introduction

Infectious diseases pose persistent threats to human health and agricultural systems and are a critical driver of wildlife population decline (Cunningham et al. 2017; Pereira et al. 2020; Capdevila et al. 2026). Disease ecology addresses these challenges by studying population-level transmission dynamics in the context of individual-level host behavior, demonstrating through growing theoretical and empirical work that host movement is central to patterns of disease spread (Altizer et al. 2011; Plowright et al. 2015; Dougherty et al. 2018; White et al. 2018; Manlove et al. 2022; Talmon et al. 2025). In recent decades, the rise of movement ecology as a prominent discipline (Nathan et al. 2008; Joo et al. 2022) has brought high-resolution animal-tracking data and new analytical tools, enabling finer-grained analyses of where infectious hosts are distributed, when infectious contacts occur, and where local outbreaks may emerge (Nathan et al. 2022; Wilber et al. 2022). Studies are also increasingly examining how variation in movement behavior across individuals, populations, and time can reshape patterns of spread, redirect epidemic trajectories, and influence the success of intervention strategies (Plowright et al. 2011; Pulliam et al. 2012; Plowright et al. 2024).

This growing interest in movement as a driver of disease dynamics has been accompanied by empirical evidence that infection can, in turn, modify movement. Sublethal haemosporidian infection in passerines can slow movement and reduce foraging ranges (Grabow et al. 2024; 2025). By contrast, gray wolves infected with toxoplasmosis are more likely to disperse from their pack (Meyer et al. 2022), and European badgers positive for bovine tuberculosis range over larger areas and forage farther from their main sett (Garnett et al. 2005). Infection can thus induce either increases or decreases in wildlife movement (Binning et al. 2017); the specific response depends on host demographics, environmental context, and the causal mechanism, which may be morphological, physiological, or neurocognitive, among others (Ezenwa et al. 2016; Talmon et al. 2025).

Although the behavioral changes we termed infection-modified movement (IMM) responses have been repeatedly documented in experimental biology and veterinary science (e.g., Hart 1988; Weary et al. 2009; Lopes et al. 2021), they are rarely incorporated into wildlife disease ecology (but see Malmberg et al. 2025; Kim et al. 2026). Traditionally, wildlife outbreak and management models treat host movement mostly as a function of environmental conditions (e.g., resource availability), conspecific density, or individual demographic traits (e.g., sex and age), but not of host disease state, thereby implicitly assuming that pre- and post-infection animal space-use patterns are epidemiologically equivalent with respect to onward transmission risk (Tao et al. 2021). This assumption likely persists for several reasons, including the scarcity of datasets that jointly capture host movement and infection dynamics, the technical difficulty of coupling these processes within a single modeling framework, and the historical separation between movement ecology, which emphasizes individual-scale behavior, from disease ecology, which operates predominately at the population scale. Nevertheless, IMM responses, which are widespread across animal taxa (Ezenwa et al. 2016; Binning et al. 2017; Talmon et al. 2025), are capable of dampening or amplifying disease spread, meaning that models that ignore them may produce biased forecasts and suboptimal management recommendations.

Modeling IMM explicitly addresses the bidirectional feedback between individual host movement and disease transmission among hosts. Although IMM responses are often localized and short-lived, confined to the interval between infection and recovery (but see post-recovery movement effects, Ingram et al. 2013), their impacts could, in principle, propagate across much broader spatial and temporal scales as infected hosts move and seed transmission elsewhere (Lopes et al. 2021).

Incorporating IMM into an infectious disease model thus provides a mechanistic basis for linking individual-scale behavioral responses with population-scale disease dynamics. The resulting theoretical framework could help assess the risk of epidemic emergence in wildlife populations, anticipate shifts in host ecology as outbreaks unfold, and inform adaptive management when real-time empirical data are limited.

Among wildlife populations, IMM may be particularly consequential in species with well-defined home ranges. Home ranging, a form of area-restricted movement driven by site fidelity, constrains both the space used by infected hosts and the number of conspecifics they may encounter while infectious, whether directly via physical contact or indirectly via exposure to viable pathogens deposited in the environment. Infection-modified home-ranging behavior could therefore modulate spatiotemporal overlap among conspecific ranges, reshaping contact structure and transmission opportunities. Home range analysis has proven valuable in this context, having been applied to predict epidemic growth (White et al. 2020), infer host connectivity (Epstein et al. 2020), and evaluate control strategies such as mass vaccination and depopulation (Cosgrove et al. 2012). Despite these contributions, home-ranging behavior has typically been modeled without feedback from infection status.

Across many species, individuals occupy home ranges not as solitary residents but as members of social groups sharing a common area (Börger et al. 2008; Powell and Mitchell 2012). We posit that both group size and the number of neighboring groups can shape disease dynamics: larger groups may accelerate intra-group transmission, whereas proximity among groups, in the absence of strong territorial avoidance, may determine how widely local outbreaks spread across the landscape. IMM should therefore be understood not only at the level of individual hosts, but also within this broader social and spatial structure, as changes in home-ranging behavior affect who infected individuals encounter as well as the degree of connectivity between infected and susceptible groups over space and time.

Here, we examine how infection-modified home-ranging behavior, represented in our model as an interplay of foraging movement and attraction to a stationary home-range center (e.g., den site), relates to final epidemic size, measured as the cumulative proportion of hosts infected over the course of an outbreak. We incorporate this movement-disease feedback into a spatially explicit individual-based model of disease spread in a wildlife population composed of multiple social groups whose home ranges can overlap. To retain generality, we focus on indirect transmission, whereby pathogens are shed into the environment and remain viable until they decay. Direct transmission arises as a limiting case when pathogen decay rates are high.

We highlight three types of IMM response: 1) anorexia, in which the infected host reduces its appetite and foraging activity, thought to serve as an immune defense that limits within-host nutrient availability and inhibits pathogen growth (Hite et al. 2019, 2020); 2) disorientation, an impairment of navigation capacity such as homing, consistent with neurocognitive dysfunction (Wolf et al. 2014); and 3) hypokinesia, a general lethargy attributable to morphological and physiological deficits (see Dekelaita et al. 2023; Nájera et al. 2025). Anorexia and disorientation act on independent components of our movement model, while hypokinesia represents their joint expression (Fig. 1); collectively, they span a broad range of possible IMM phenotypes. In this work, we focus on these symptomatic responses that modify individual space-use pattern, setting aside movement strategies aimed at exploiting, managing, or avoiding infection, whether parasite-driven, as in parasitic manipulation (Stockmaier et al. 2021), or host-driven, as in migratory recovery (Shaw and Binning 2016) and migratory escape (Altizer et al. 2011). Our model also does not address higher-order effects, such as changes in the social interactions or group behavior of uninfected individuals in response to local infections (Stockmaier et al. 2021).

**Figure 1.**
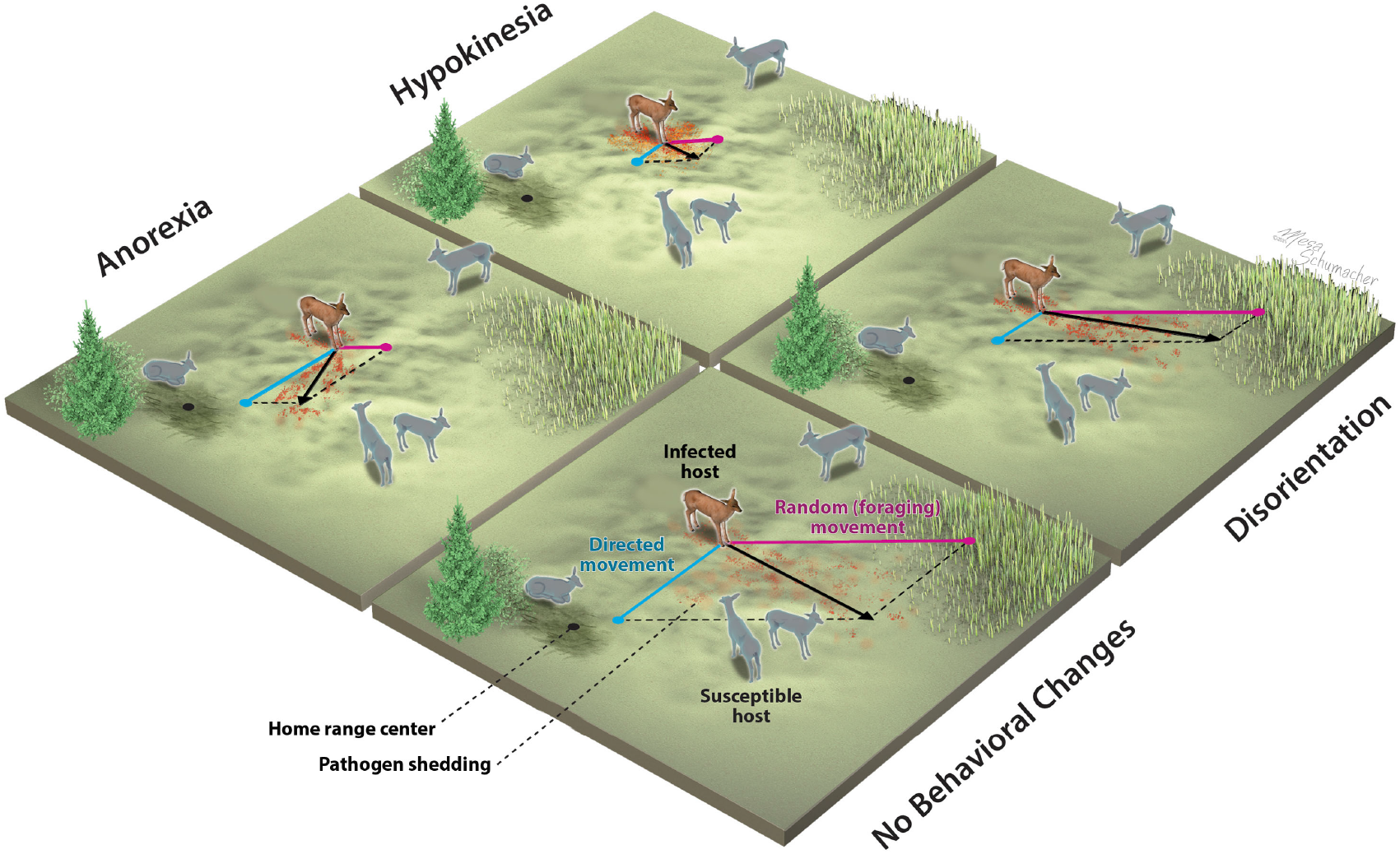
Three types of infection-modified movement responses (IMM) compared to the conventional modeling approach that assumes no infection-induced behavioral changes. Each response is defined by modifications to specific components of the Ornstein-Uhlenbeck equation describing an individual host’s area-restricted movement: anorexia reduces the directionally random (diffusive) component associated with foraging, disorientation reduces the directed (drift) component biased toward a stationary home-range center, and hypokinesia reduces both components equally. These movement responses influence the spatiotemporal distribution of shed pathogens and, by extension, transmission risk to susceptible hosts. Illustration credit: Mesa Studios (www.mesaschumacher.com).

The effects of IMM likely depend on several fundamental features of wildlife movement and disease ecology: host group structure, baseline home-ranging behavior, and the duration of pathogen persistence in the environment. Each may vary seasonally in natural wildlife systems: social grouping in many species undergoes seasonal fission-fusion dynamics (Wittemyer et al. 2005; Webber and Vander Wal 2021); home range size and space-use pattern typically track seasonal resource distribution and life-history demands (e.g., mate search, reproduction) (Börger et al. 2008; van Beest et al. 2011); and pathogen viability in environmental reservoirs is often sensitive to temperature and humidity, with some pathogens decaying more slowly during fall/winter season (e.g., *Mycobacterium bovis*, Fine et al. 2011). To capture these disparate sources of seasonality, we further simulate independent fluctuations in each factor and assess how they modulate the epidemic role of IMM.

In wildlife species with clear sociality, conspecific interactions, whether expressed as spatial attraction or avoidance, can contribute substantially to disease transmission and spread (White et al. 2018; Yang et al. 2021; Gilbertson et al. 2023). Like IMM, conspecific interactions enable restructuring of host contact networks, therefore, we posit that these two processes may compound or offset one another in shaping disease outcomes. As a second model extension, we investigate their combined epidemic effects by introducing the phenomenon of social cohesion, defined as mutual attraction of group members during individual movement.

Our spatially explicit, individual-based simulation models capture a dynamic interdependence between host movement and disease transmission within wildlife populations. They constitute a new theoretical framework that integrates classical modeling elements from wildlife disease ecology and movement ecology. We simulate epidemics with and without IMM, comparing predicted outcomes to assess the general need to account for movement-disease feedback in future wildlife ecology research.

## Results

Infected individuals can occupy home ranges much larger or smaller than those of susceptible individuals, depending on which movement component is modified: foraging (diffusion), site-fidelity (advection), or both (Fig. 2a). Under infection-induced anorexia, home ranges contract, reflecting reduced diffusive movement associated with foraging activities. Under infection-induced disorientation, site-fidelity weakens and home ranges expand. When infection induces hypokinesia by attenuating both components equally, home ranges contract as under anorexia, though less sharply. Home range size is approximately preserved when the advection component is reduced by the square of the factor scaling the diffusion component, consistent with analytical predictions derived from the classical Fokker-Planck equation modeling animal space use (Moorcroft and Lewis 2013; Tao et al. 2025).

**Figure 2.**
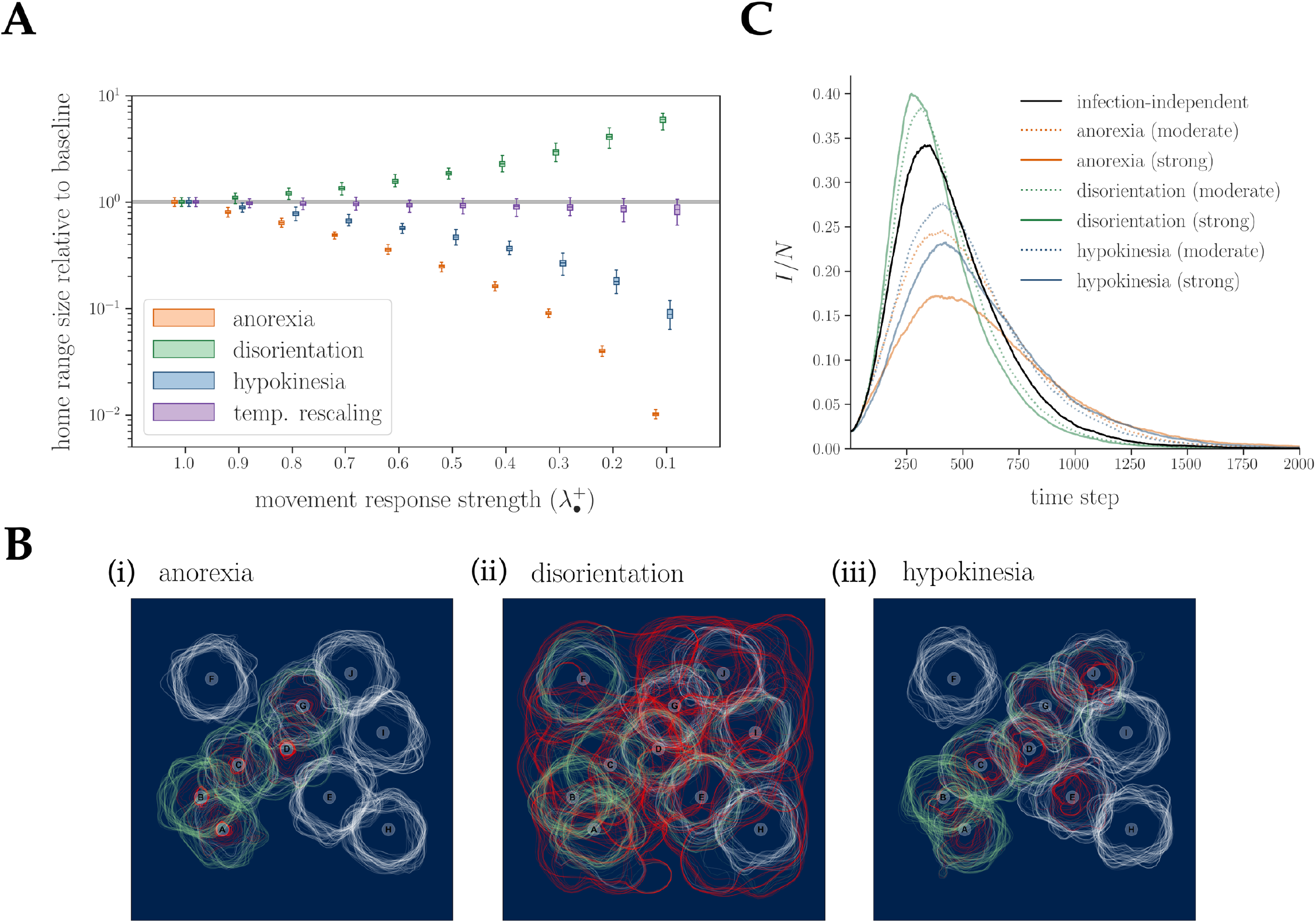
Effects of infection-modified movement responses on host home range size, space-use overlap, and disease dynamics. (a) Box plots show how home range size deviates from the infection-independent baseline under four IMM responses: anorexia, disorientation, hypokinesia, and temporal rescaling. In each, 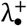 denotes the varied coefficient: 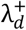 (anorexia) and 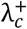 (disorientation), with the other fixed at 1, or 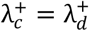 (hypokinesia). For temporal rescaling, 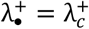 with 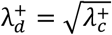, leaving home range roughly invariant. Home range sizes were estimated as the area within the 90% utilization distribution contour, computed from 2,000 relocations per replicate (100 replicates per response) using the adehabitatHR package (version 0.4.21) in R, and are expressed as a fraction of the baseline median. The grey band marks the baseline interquartile range (100 replicates). (b) Space-use patterns of ten neighboring individuals (A-J) with independent home-range centers, assuming infection-induced anorexia 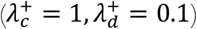, disorientation 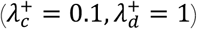, or hypokinesia 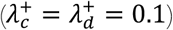. Each line denotes a host’s 90% utilization distribution contour, estimated from movement within a sliding 200-step window (i.e., steps 0-200, 15-215, 30-230, …) over a total of 1,500 time steps, colored by majority infection status during the corresponding window (white, susceptible; red, infected; green, recovered). (c) Mean fraction infected (*I*/*N*) over time under selected IMM responses, averaged across 100 replicates. The host population (*N* = 50) occupies a landscape with periodic boundaries. Home-range centers are individually randomized (no group formation). Moderate and strong responses correspond to reductions in the relevant coefficient(s) to 0.5 and 0.1, respectively. In (b-c), illustrative parameters: pathogen decay rate *δ* = 0.05; baseline diffusion coefficient *σ* = 3; baseline drift coefficient *θ* = 0.2; infection rate *β* = 0.2; recovery rate *γ* = 0.05.

In our simulations, space use fluctuates naturally due to the stochasticity of the underlying movement process, with additional, potentially larger fluctuation introduced when a host transitions between different disease states. These infection-driven fluctuations can intensify the use of core areas and accelerate local environmental pathogen buildup, or generate transient space-use overlap that increases the frequency of previously rare encounters (Fig. 2b; Supplementary Video 1).

The type and strength of the IMM response therefore determine epidemic trajectories. This effect stems from response-specific changes in a host’s home-range size during infection: contractions under anorexia and hypokinesia; expansions under disorientation (Fig. 2a). These space-use dynamics reorganize the host’s direct and indirect contacts, altering the speed and extent of disease spread (Fig. 2b). Consequently, even in the simplest case, where each host is its own group and moves around an independently located home-range center (group number *m* = population size *N*), incorporating IMM responses can visibly shift the epidemic curve (Fig. 2c): in these illustrative runs, anorexia and hypokinesia lower and delay the epidemic peak while sustaining high incidence over a longer term, whereas disorientation drives an earlier, higher peak that declines more quickly.

The relative epidemic response (RER) measures the epidemic impact of IMM responses relative to the conventional assumption of infection-independent movement (Fig. 3a-b). In single-group populations (*m* = 1), anorexia clustered infected hosts near the shared home-range center and increased mean final epidemic size ⟨ℛ⟩ from 0.95 to as high as 1. Disorientation, by contrast, dispersed infected individuals away from susceptible conspecifics and reduced ⟨ℛ⟩ from 0.95 to as low as 0.75. The effect of hypokinesia was negligible. Across IMM responses, ⟨ℛ⟩ varied by 20 percentage points. (Fig. 3c).

**Figure 3.**
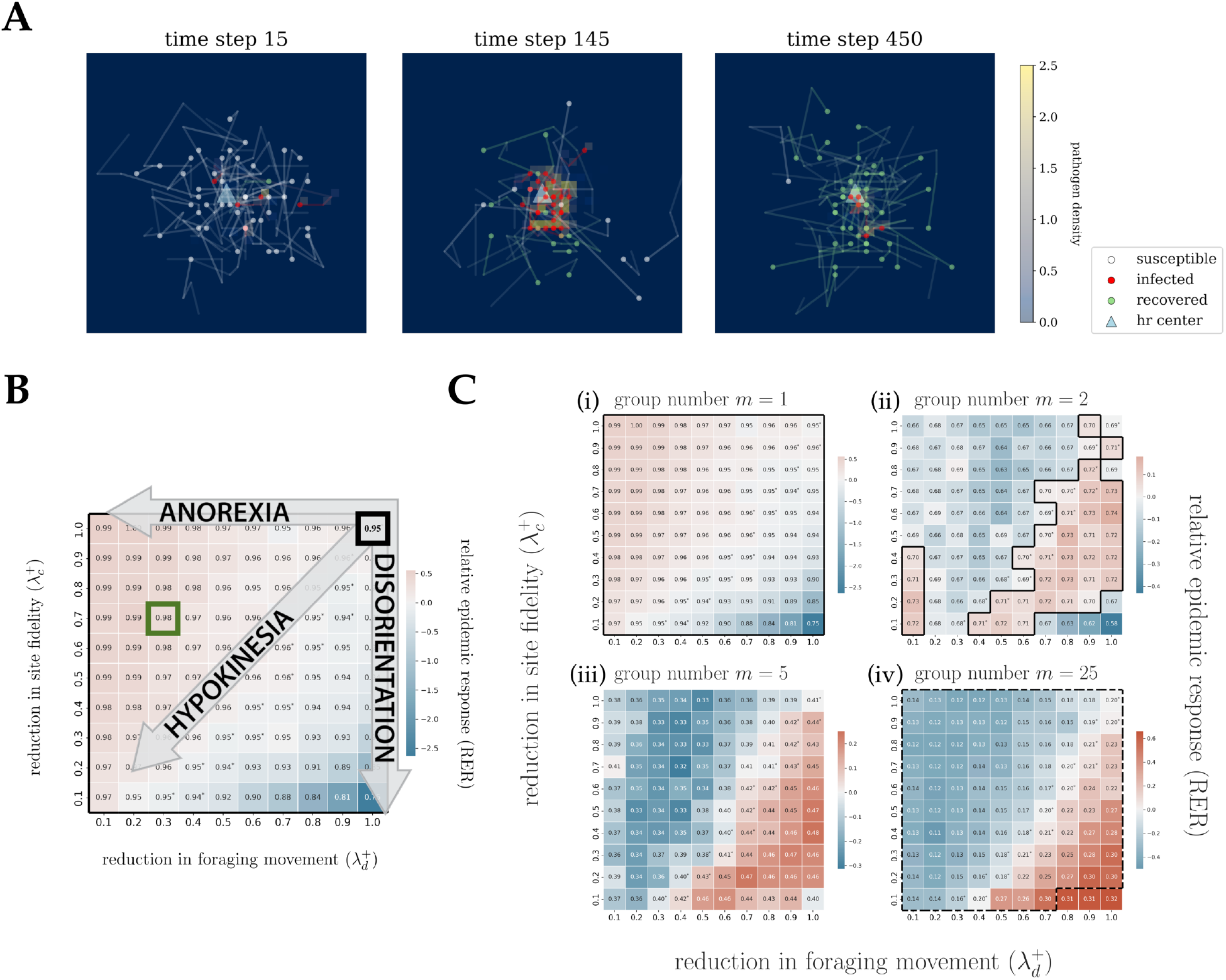
(a) Snapshots from an example individual-based simulation of disease spread in a single-group population (*N* = 50) with an IMM response defined by 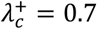 and 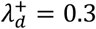 (outlined in green in panel b). (b) Schematic for interpreting panel c, illustrated using the grid for group number *m* = 1; cell colors and values are defined in (c). The cell in the upper-right corner (outlined in black) represents the epidemic outcome under infection-independent movement 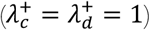 and serves as the reference. Arrows in IMM parameter space indicate increasing strength of anorexia, disorientation, and hypokinesia. (c) Relative epidemic response (RER; equations 2a-b) across IMM parameter combinations under varying grouping structure (group number *m* = 1, 2, 5, and 25; total population size *N* = 50 held constant, so group size equals *N*/*m*). Red indicates positive RER (larger epidemics relative to the reference); blue indicates negative RER (smaller epidemics). Each cell shows ⟨ℛ⟩, the mean final epidemic size for that parameter combination across 750 simulation replicates. Solid contours enclose ⟨ℛ⟩ ≥ 0.70; dashed contours enclose ⟨ℛ⟩ ≤ 0.30. Asterisks mark cells where infected and uninfected individuals have home ranges of roughly comparable area 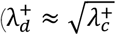; “temporal rescaling” in Fig. 2a). Pathogen decay rate *δ* = 0.25; baseline diffusion coefficient *σ* = 3; baseline drift coefficient *θ* = 0.2; infection rate *β* = 0.1; recovery rate *γ* = 0.005.

When we modeled disease dynamics in populations of different grouping structures, identical IMM responses produced contrasting epidemic outcomes. In fragmented, multi-group populations (*m* > 1), overall ⟨ℛ⟩ declined with the number of groups (Fig. 3c). There, anorexia appears to promote group self-isolation, constraining transmission between groups. The standardized Morisita’s indices *I*_*p*_ (Smith-Gill 1975) are consistent with this mechanism, showing that many anorexia-like IMM responses 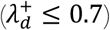 concentrated infections within subsets of groups rather than distributing them randomly across the population (Supplementary Fig. 1). Disorientation, in contrast, seems to enhance transmission by enabling infected hosts to reach otherwise unexposed groups through longer-distance movement (Figs. 3c). In highly fragmented populations (*m* = 25), maximal disorientation increased ⟨ℛ⟩ by 0.12 (Fig. 3c). Similar reversals arose even among IMM responses that left home range size largely unchanged (Fig. 3c, cells marked with asterisks). Hypokinesia became increasingly epidemic-suppressing with fragmentation (Fig. 3c).

Varying environmental pathogen decay (*δ*) and the baseline diffusion of uninfected hosts (*σ*) reshaped both the magnitude and direction of IMM effects on epidemics (Fig. 4). Higher *δ* and *σ* generally reduced ⟨ℛ⟩ while widening its range across IMM responses: in single-group populations (*m* = 1), ⟨ℛ⟩ ranged from 0.20 under maximal disorientation to 0.80 under maximal anorexia, a 60 percentage-point spread attributable to movement-disease feedback alone. Hypokinesia exerted a notable epidemic-enhancing effect, raising ⟨ℛ⟩ from 0.39 to as high as 0.54. The directional reversal of anorexia effects previously observed at *m* = 2 (Fig. 3c) did not emerge until *m* = 25 (Fig. 4); in these highly fragmented populations, disorientation’s epidemic-enhancing effect largely dissipated, while hypokinesia remained epidemic-suppressing. Standardized Morisita’s indices stayed near zero across IMM responses (Supplementary Fig. 2). Overall, these results demonstrate that the spatial signature and epidemic consequences of IMM responses depend fundamentally on ecological context.

**Figure 4.**
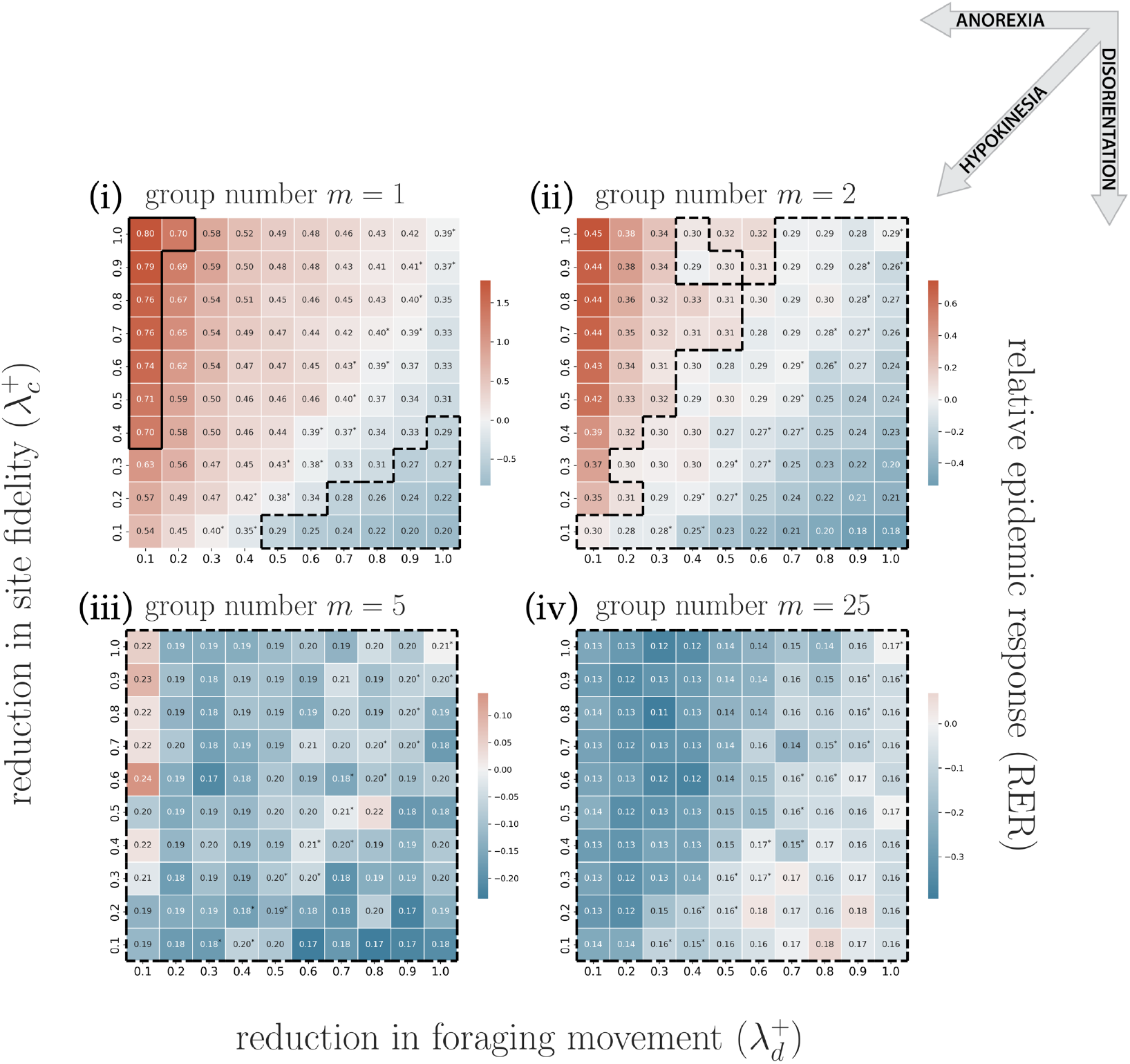
Relative epidemic response (RER) across IMM parameter combinations under varying grouping structure (group number *m* = 1, 2, 5, and 25), with pathogen decay rate *δ* = 0.35 and baseline diffusion coefficient *σ* = 6. All other parameters and plotting conventions as in Fig. 3c.

Sampling ecological conditions from different phases of the seasonal cycle across multiple years, we found that IMM responses drove systematic deviations from the conventional no-IMM predictions (Fig. 5). Anorexia tended to increase ℛ, with deviations apparent at three of the four sampled time points (*t*_0_, *t*_1_, *t*_2_), reaching nearly 0.9 at *t*_1_, when individuals were partitioned into few groups, foraged extensively, and pathogen decay was moderate (Fig. 5a). In contrast, disorientation generally reduced ℛ from *t*_0_, *t*_1_, and *t*_2_, most strikingly at *t*_2_, when group number was high, foraging activity was low, and pathogen decay was moderate; here, outbreaks often failed to establish, as opposed to the large epidemics (ℛ ≈ 0.8) predicted by the no-IMM model (Fig. 5b). At *t*_3_, when hosts were maximally fragmented and sedentary, and pathogens decayed rapidly, predictions from both IMM responses converged most closely with the no-IMM model, though anorexia yielded slightly smaller and disorientation slightly larger epidemics (Fig. 5a-b). Therefore, the conventional, no-IMM model over- or underestimated epidemic outcomes relative to those under IMM according to the seasonal condition, with the magnitude of bias ranging from modest to substantial.

**Figure 5.**
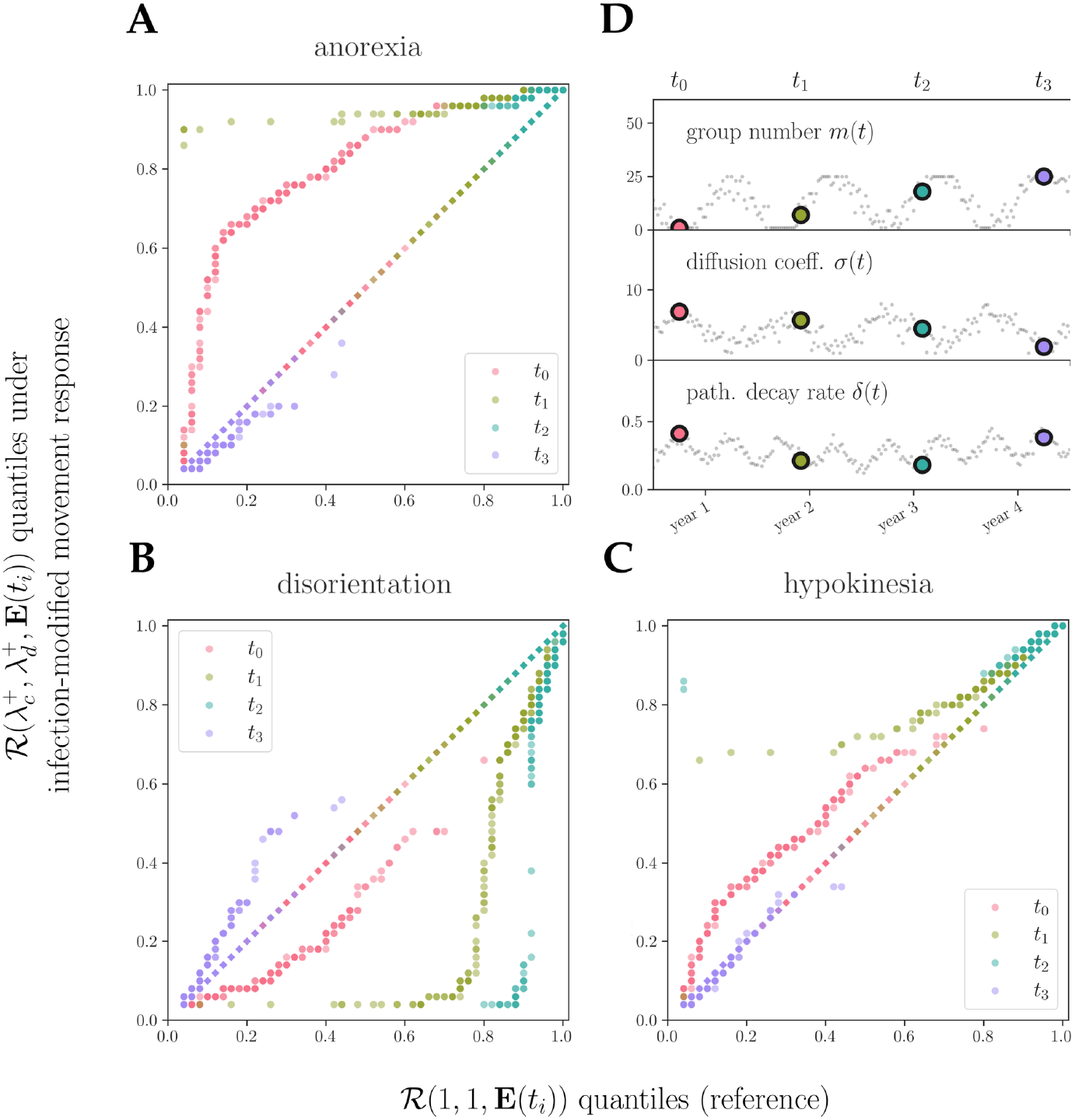
Epidemic outcomes under multi-year cycles of seasonally varying ecological conditions **E**(*t*_*i*_ = {*m*(*t*), *σ*(*t*), *δ*(*t*)}. (a-c) Quantile-quantile plots comparing the distribution of final epidemic sizes under three IMM responses, 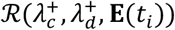, against the no-IMM (infection-independent movement) reference, ℛ(1,1, **E**(*t*_*i*_)), each distribution computed from 200 simulation replicates per time point *t*_*i*_ = *t*_0_, …, *t*_3_ as colored in (d): (a) anorexia 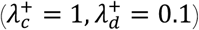, (b) disorientation 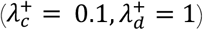, and (c) hypokinesia 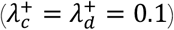. The diagonal is the no-IMM reference plotted against itself; points above and below it indicate epidemic-enhancing and epidemic-suppressing effects, respectively. (d) Seasonal variation in group number *m*(*t*), baseline diffusion coefficient *σ*(*t*), and pathogen decay rate *δ*(*t*), with colored circles marking parameter values at evenly spaced times: **E**(*t*_0_) = {1, 6.88, 0.41}, **E**(*t*_1_) = {7, 5.67, 0.21}, **E**(*t*_2_) = {18, 4.48, 0.18}, **E**(*t*_3_) = {25, 1.89, 0.38}. All remaining parameters are held fixed as in Fig. 3c.

Hypokinesia produced a different pattern. ℛ values either exceeded (*t*_0_, *t*_1_) or closely followed (*t*_2_, *t*_3_) the no-IMM predictions (Fig. 5c). At *t*_2_, the two models nearly coincided across the upper quantiles, both predicting large outbreaks, but hypokinesia raised the lower tail: its smallest ℛ values were far larger than the smallest under no-IMM, so more epidemics successfully took hold.

Hypokinesia may therefore meaningfully impact epidemic dynamics by occasionally buffering against stochastic fade-out, despite having smaller effects on ⟨ℛ⟩ than anorexia or disorientation (Figs. 3-4).

Introducing social cohesion into host movement allows infected individuals to remain near their groupmates in predominantly uninfected groups, counteracting the movement-disease feedback that would otherwise separate them from the group (Fig. 6; Supplementary Fig. 3). When host diffusivity is high (*σ* = 6), increasing cohesion strength *κ* generally elevated ⟨ℛ⟩ while narrowing its range across IMM responses. Stronger cohesion also lessened or reversed the epidemic effect of IMM across most of the parameter grid, notably at high 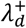. In two-group populations (*m* = 2), for example, a large class of IMM responses that suppressed transmission under no cohesion enhanced it under strong cohesion: at 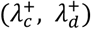 = (0.1, 0.8_*i*_, varying *κ* from 0 to 1 moved ⟨ℛ⟩ from below to above the infection-independent reference (0.49 vs. 0.57, then 0.72 vs. 0.70; Fig. 6). A similar reversal occurred in more fragmented populations (*m* = 5), where cohesion further concentrated infection within groups: under strong anorexic or hypokinetic responses, infected individuals were significantly aggregated by group identity (Fig. 6; Supplementary Fig. 4).

**Figure 6.**
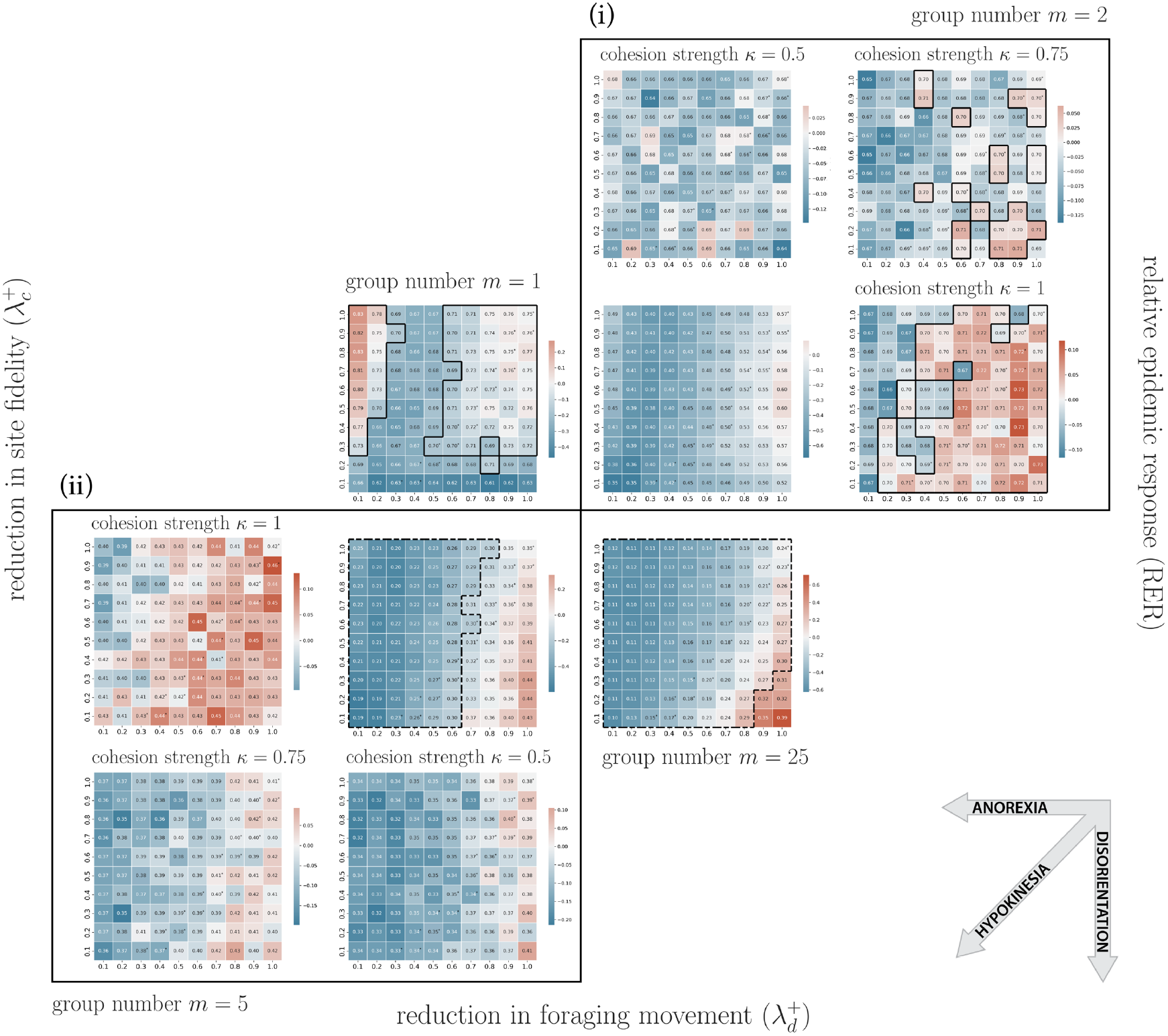
Effect of social cohesion on relative epidemic response (RER). The four central heatmaps show the *κ* = 0 baseline scenario. Additional panels surrounding the *m* = 2 (top-right) and *m* = 5 (bottom-left) heatmaps correspond to cohesion strengths *κ* = 0.5, 0.75, and 1, arranged clockwise. Landscape boundaries are hard rather than periodic (cf. Figs. 3 and 4). All other parameters and plotting conventions as in Fig. 4.

## Discussion

We introduced a modeling framework that formalizes a reciprocal relationship commonly observed in wildlife systems: the movement of individual hosts following infection determines pathogen exposure risk for conspecifics, generating new infections that in turn modify the movement of other individuals. By extending a traditional spatial disease modeling approach with infection-modified movement (IMM) responses, we simultaneously addressed two forms of dynamical feedback: between movement and disease transmission, and between individual- and population-scale processes.

We showed that prototypical IMM responses, exemplified by anorexia, disorientation, and hypokinesia, can drive divergent epidemic outcomes, and that these effects reverse direction as grouping structure changes. The impact of IMM responses further depends on environmental attributes, including conditions affecting pathogen decay and baseline home-ranging behavior. Social behavior adds another layer of complexity: social cohesion can override the spatial separation that an IMM response would otherwise induce between infected and susceptible hosts, sustaining transmission opportunities and thereby lessening or reversing its epidemic effect. Across seasonal cycles, conventional models applied to systems where IMM has been documented may therefore oscillate between over- and under-estimation of epidemic severity as the ecological context shifts over time. Disease forecasts that treat host movement as infection-independent will systematically overlook the dynamics generated by such movement-disease feedback.

Changes in individual movement and social contact are routinely incorporated into models of human diseases and their management, both as self-initiated behavioral responses (Perra et al. 2011; Verelst et al. 2016) and policy interventions (Ferguson et al. 2005; Flaxman et al. 2020), but studies of wildlife disease still largely treat host movement as fixed. In our model, infection systematically alters short-term wildlife movement at the individual scale; altered movement then propagates through the population alongside the pathogen, reshaping long-term epidemic trajectories. In line with recent proposals to base wildlife disease surveillance on detection of movement anomalies (Talmon et al. 2025), our results suggest that mass movement monitoring may help monitor epidemic potential in real time for populations that are difficult to sample directly. As tracking technologies become more accessible, this approach is particularly suited to zoonoses and livestock diseases with cryptic wildlife reservoirs, such as Rift Valley fever, where serological surveys are often slow and logistically constrained (Rostal et al. 2025).

The spatial structure of infection in our simulations informs intervention design. Anorexia, for example, can concentrate the risk of pathogen exposure in high-traffic areas around the home-range center, creating infection hotspots amenable to targeted intervention. More broadly, when infections are statistically aggregated within a subset of groups, strategies such as vaccination or culling of identified groups become more practical than when infections are widely distributed across the population. Because aggregation patterns vary with IMM responses and ecological conditions, effective management will require matching intervention strategies to the underlying behavioral and environmental processes. Methods for optimizing such targeted interventions under logistical constraints are developed in Tao et al. (2018, 2021b).

Adding habitat fragmentation to our spatially homogeneous model could reconfigure host-host and host-pathogen interactions through several mechanisms. Loss of local resources can expand host foraging ranges or relocate home-range centers, both promoting between-group transmission; higher edge-to-interior ratios can reshape local temperature and humidity, which govern environmental pathogen persistence; and greater group isolation can restructure host contact networks at regional scales. Given the global intensification of habitat fragmentation, modeling movement-disease feedback in spatially explicit, fragmented landscapes could enable epidemic forecasts that capture behavioral and spatial complexity within a metapopulation framework, and could challenge classical assumptions in wildlife disease ecology (for a related approach, see Tao et al. 2024).

Infection can be viewed as a transient phase in host behavior: the individual deviates from its typical movement pattern while infected and returns to it (or not) upon recovery. Transient dynamics have recently gained traction in disease ecology and epidemiology, including in the spatial dynamics driven by individual host movement (Daversa et al. 2017; Tao et al. 2021a). This extends a longer-standing recognition of transients in ecological systems in general (Hastings 2004; Hastings et al. 2018; Morozov et al. 2020). By formalizing IMM, we linked the duration and expression of this transient phase to epidemic outcomes. Two recent methods provide the tools to advance transient analysis at the interface of movement and disease ecology. Tao et al. (2025) introduced a computational framework that solves Fokker-Planck equations beyond their classical steady-state solutions, resolving the transient phase of area-restricted space use; the same approach could be applied to a system in which these equations are coupled to spatial compartmental models, characterizing how host space-use patterns and disease states evolve jointly over time. In addition, recently developed movement-driven modeling of spatiotemporal infection risk (MoveSTIR) leverages observed movement trajectories to predict transmission dynamics on real-world landscapes (Wilber et al. 2022; Vargas Soto et al. 2025). The transient effects of IMM responses on disease dynamics could be explored retrospectively by perturbing the trajectory segments that correspond to sickness behavior. This would reveal how sickness duration and the resulting transient utilization distributions influence each host’s contribution to transmission risk.

A key benefit of our approach is its testability: variants of the Ornstein-Uhlenbeck model underlying our framework are routinely fit to wildlife tracking data (e.g., Hooten et al. 2017), which are increasingly collected to aid wildlife disease surveillance (Pepin et al. 2025). When both longitudinal movement and infection data are available for a representative sample of individuals (Giglio et al. 2025), one can fit these models to the movement data with time-varying infection status as a covariate, and test how the drift and diffusion coefficients change with it. In many populations, however, longitudinal infection data remain difficult to collect. Hidden Markov models (Kim et al. 2026) and deep learning (Blaha et al. 2026) can overcome this limitation, inferring IMM responses directly from movement data without requiring infection data. By estimating how each coefficient changes with inferred infection status for a given host and pathogen, these methods could supply the parameters our framework needs to establish movement-disease feedback as a measurable property of wildlife disease systems, rather than leaving it an unquantified source of bias.

Our model defines IMM responses as reductions in one or two basic movement components that comprise simple home-ranging behavior 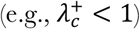, yet other forms of response have been documented. For example, a compensatory increase in movement, where infected hosts forage more to meet elevated metabolic costs of immune activation (Povey et al. 2009), could map onto the same framework with 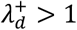. Infection can also induce directional preferences toward landmarks such as water sources when extreme thirst drives resource selection (Barrile et al. 2024) or reduce hosts’ ability to discriminate among habitats of differing quality (Grabow et al. 2024). Apart from the behavior of sick individuals, infection can influence the movement of healthy group members through active caregivng, observed most clearly in eusocial systems (Stockmaier et al. 2021). Finally, inter-group avoidance through strict territoriality can suppress epidemic severity while prolonging persistence (White et al. 2020); the effects of infection-dependent territoriality, however, remain unexplored.

In addition to direct changes to movement behavior, infection may alter how hosts acquire, process, and store spatial information (Townsend et al. 2022). Such cognitive impairment could shape individual-level movement decisions and, in turn, population-level disease dynamics. As rapid environmental change degrades the reliability of fitness-relevant sensory cues, infection-driven cognitive processes could disrupt space-use patterns and contact networks, generating novel opportunities for onward transmission. Extending our framework with tools from information theory (Stevens 2013; Bergman and Beehner 2023) would bridge sensory ecology with movement and disease ecology, thereby opening a new avenue for understanding epidemic risk in the face of increasing environmental uncertainty.

## Methods

We modeled the system with individual-based simulations on a homogeneous two-dimensional landscape, represented as a 100 × 100 square lattice with periodic boundary condition to eliminate edge effects. The population consisted of functionally identical host individuals (no demographic structure). To isolate the effects of movement-disease feedback from population dynamics, we kept population size constant by excluding mortality, reproduction, and immigration (emigration) from the model.

Each host *i* moves independently according to an Ornstein-Uhlenbeck process describing a continuous-time biased random walk indexed by time *t*. This movement process comprises two components: a) isotropic Brownian diffusion *W*_*i,t*_, a Wiener process representing foraging, and b) drift toward the host’s stationary home-range center, representing site fidelity. The home-range center is defined by a point attractor, 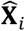, and the drift term is mean reverting, with attraction toward 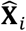 increasing with distance from the center. The host’s location, **X**_*i,t*_, evolves as follows:

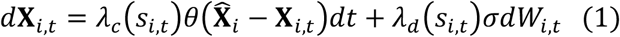

The parameters *θ* and *σ* denote the baseline drift and diffusion coefficients, respectively; together, they determine the host’s utilization distribution, with stronger drift relative to diffusion producing a smaller, more intensively used home range.

The disease portion of our model follows the canonical Susceptible-Infected-Recovered (SIR) framework, with the additional assumption that the infection may temporarily modify host movement upon transition from the susceptible to the infected state (*S* → *I*) by scaling either the foraging component, the site-fidelity component, or both, through *λ*_*c*_(*s*_*i,t*_ = *I*) and *λ*_*d*_(*s*_*i,t*_ = *I*), where *s*_*i,t*_ ∈ {*S, I, R*} denotes the disease state of host *i* at time *t*. For brevity, we denote the infected-state values of these parameters hereafter as 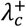 and 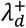. Upon recovery (*I* → *R*), these pathogenic effects cease and both the foraging and site-fidelity components return to baseline values, with *λ*_*c*_(*S*) = *λ*_*c*_(*R*) = 1 and *λ*_*d*_ (*S*_*i*_) = *λ*_*d*_ (*R*) = 1.

We considered three forms of infection-modified movement (IMM): anorexia, a potential immune response that reduces foraging activity and thus limits diffusive movement 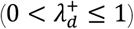; disorientation, an impairment in homing ability that weakens site fidelity 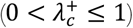; and hypokinesia, a general reduction in locomotive capacity 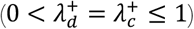.

We discretized the infection-dependent Ornstein-Uhlenbeck equation (Eq. 1) and simulated host movement trajectories using the Euler-Maruyama method (Maruyama 1955; Guo et al. 2018). For model tractability, we excluded from the base model both direct behavioral interactions, including conspecific attraction and avoidance, and indirect behavioral interactions arising through space or resource competition.

Transmissions occur indirectly through an environmental reservoir. At each time step (Δ*t* = 1, without loss of generality), each infected host shed one unit of pathogen into the lattice cell it occupies, allowing pathogen to accumulate locally through repeated visitation and shedding.

Environmental pathogens decay geometrically at rate *δ*, such that the contribution of pathogens deposited at time *t*, is discounted by a factor of 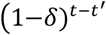 at time *t*. The resulting environmental pathogen load at location **x** and time *t* is therefore given by 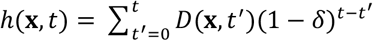, where *D*(**x**, *t*^’^) denotes the quantity of pathogens deposited at location **x** at time *t*,. A susceptible host entering a lattice cell containing viable pathogens at time *t* becomes infected with probability 1 ™ exp[™*βh*(**x**, *t*)], where *β* is the infection rate. Infected hosts recover at rate *γ* and acquire lifelong immunity, precluding reinfection.

We initialized each simulation with *N* = 50 individuals evenly distributed among *m* groups (*m* ≤ *N*). Each group shared a common home-range center that was randomly located on the landscape at the start of each simulation and then held fixed over time. We assumed that stable home ranges were established prior to pathogen introduction. Accordingly, we ran the model with susceptible hosts alone for a burn-in period of 1,000 time steps before infecting two randomly chosen individuals. We then continued the simulation for an additional 2,500 time steps.

Host movements and pathogen transmissions were then simulated jointly until no infected hosts remained, after which we recorded the final epidemic size, ℛ, defined as the total fraction of hosts that were infected. For each IMM response parameterized by 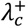 and 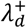, we performed *n*_rep_ simulation replicates. To quantify the impact of IMM, we computed a *z*-score for each replicate *k* relative to the reference case 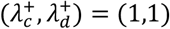:

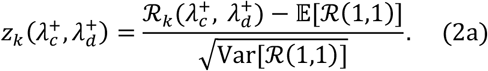

We then averaged *z*-scores across replicates,

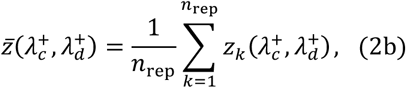

which we term relative epidemic response (RER), a dimensionless measure of the extent to which epidemic outcomes under a given IMM response diverged from those under the conventional assumption of infection-independent movement.

To characterize how IMM responses shape epidemic outcomes, we computed RER across a grid of 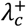 and 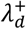 values. We first examined how group number *m* shapes the resulting profile. We then repeated this analysis under different values of *δ* and *σ* to assess how a broader set of ecological conditions modulates these profiles. We also computed the standardized Morisita’s dispersion index *I*_*p*_ on the number of infections in each group using the ‘vegan’ package (version 2.6-4) in R, then averaged across replicates, to quantify the social aggregation of infection by group identity.

Next, we examined seasonal effects by allowing *m, σ*, and *δ* to all vary periodically over time, collectively denoted **E**(*t*) = {*m*(*t*), *σ*(*t*), *δ*(*t*)}, representing seasonal forcing. We constructed **E**(*t*) to capture observed features of seasonality in host and pathogen: a) *m*(*t*) is bounded, with each bound held for part of the year to represent sustained periods of strong mixing and strong fragmentation, as seen in the social dynamics of many cervids, (e.g., DeYoung and Miller 2011); b) *m*(*t*) and *σ*(*t*) follow annual cycles in antiphase (see Ofstad et al. 2016); c) host movement is sensitive to stochastic resource availability (Singer et al. 1981), which we represent by giving *σ*(*t*_*i*_ the lowest signal-to-noise ratio of the three; d) *δ*(*t*) follows a sub-annual cycle, which we set to semiannual, tracking variation in temperature and humidity (Morris et al. 2021); e) *δ*(*t*)’s annual mean drifts between years by an amount comparable to its semiannual amplitude, a proxy for interannual variation in those conditions. We assumed that the timescale of any outbreak initiated at time *t*_*i*_ is short relative to seasonal change in these parameters, such that the forcing remains approximately constant at **E**(*t*_*i*_) over its duration. To test whether and how seasonality may affect the importance of IMM-based epidemic predictions, we compared the distribution of ℛ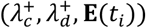 across simulation replicates against that of ℛ(1,1, **E**(*t*_*i*_)), for equally spaced times *t*_0_, …, *t*_3_, using quantile-quantile plots.

Finally, we relaxed the assumption of independent host movement to investigate the effects of IMM under different levels of social cohesion, which we defined as the tendency for members of the same group to remain spatially proximate during movement. To incorporate this additional behavior, we extended the infection-dependent Ornstein-Uhlenbeck equation (Eq. 1) to include attraction toward the current centroid of the focal individual’s group,

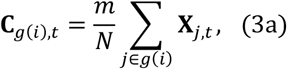

such that

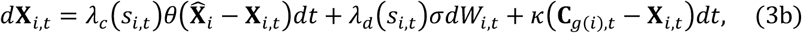

where *g*(*i*) denotes the group containing individual *i*, and *κ* ≥ 0 controls the strength of social cohesion. For this analysis, we imposed a hard (fixed) boundary condition on the landscape to avoid artificial toroidal (wraparound) interactions among spatially distant individuals that could otherwise mask differences between weakly and strongly cohesive groups.

The model simulations were implemented in MATLAB (version R2023a). To improve computational efficiency, we used vectorized operations and parallelized simulations with MATLAB’s Parallel Computing Toolbox, while offloading core computations to the GPU using gpuArray.

## Acknowledgement

We thank Paul Blackwell and Valeria Giunta for their technical input.

