## Supplementary Figures for "Neglecting movement-disease feedback biases wildlife epidemic forecasts"

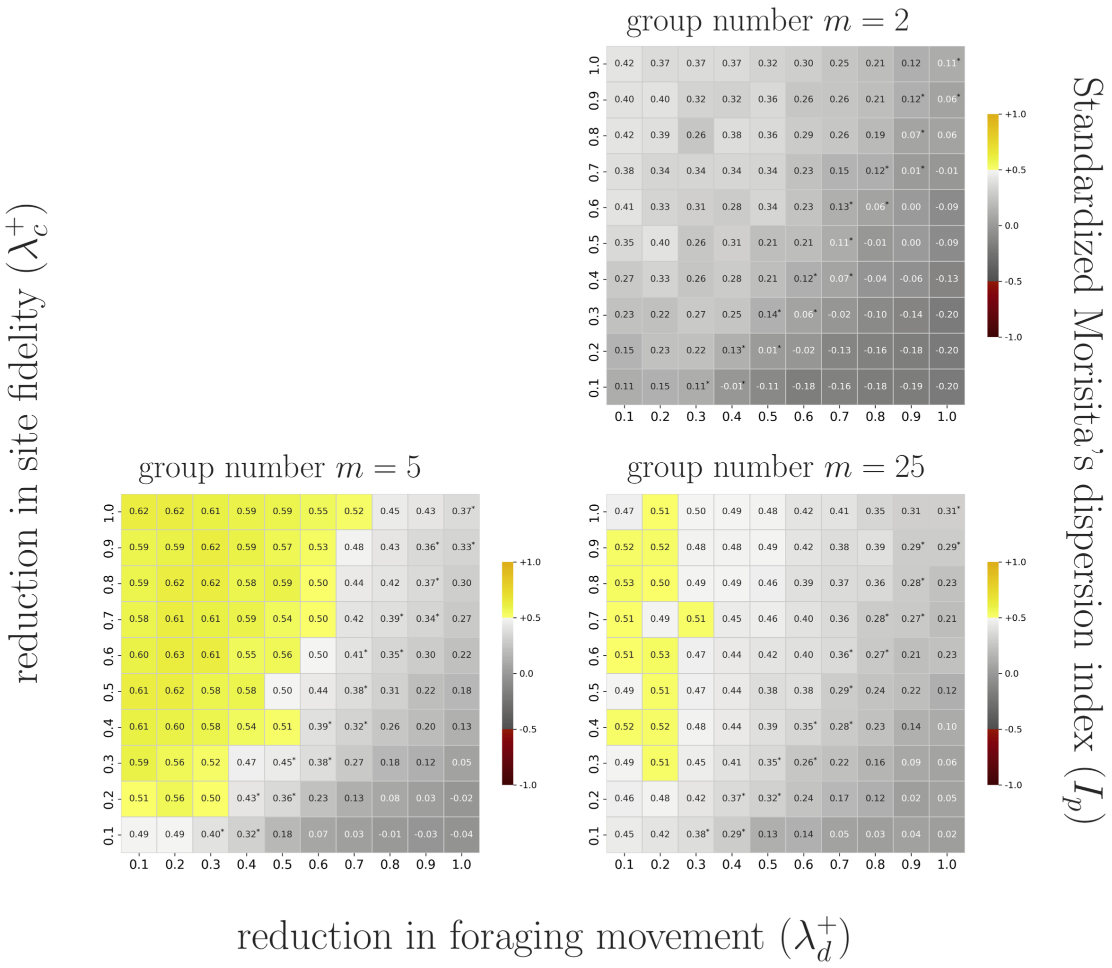


Supplementary Figure 1. Standardized Morisita's dispersion index $I_p$ measuring the aggregation of infection among groups, across infection-modified movement (IMM) parameter combinations and three grouping structures (group number $m = 2, 5,$ and $25$). Each panel maps directly onto its counterpart in Fig. 3c.ii-iv, omitting the $m = 1$ case (Fig. 3c.i) where the index is undefined. $0.5 \leq I_p \leq 1$ (yellow) indicates that infections are concentrated in a subset of groups. All parameters as in Fig. 3c.

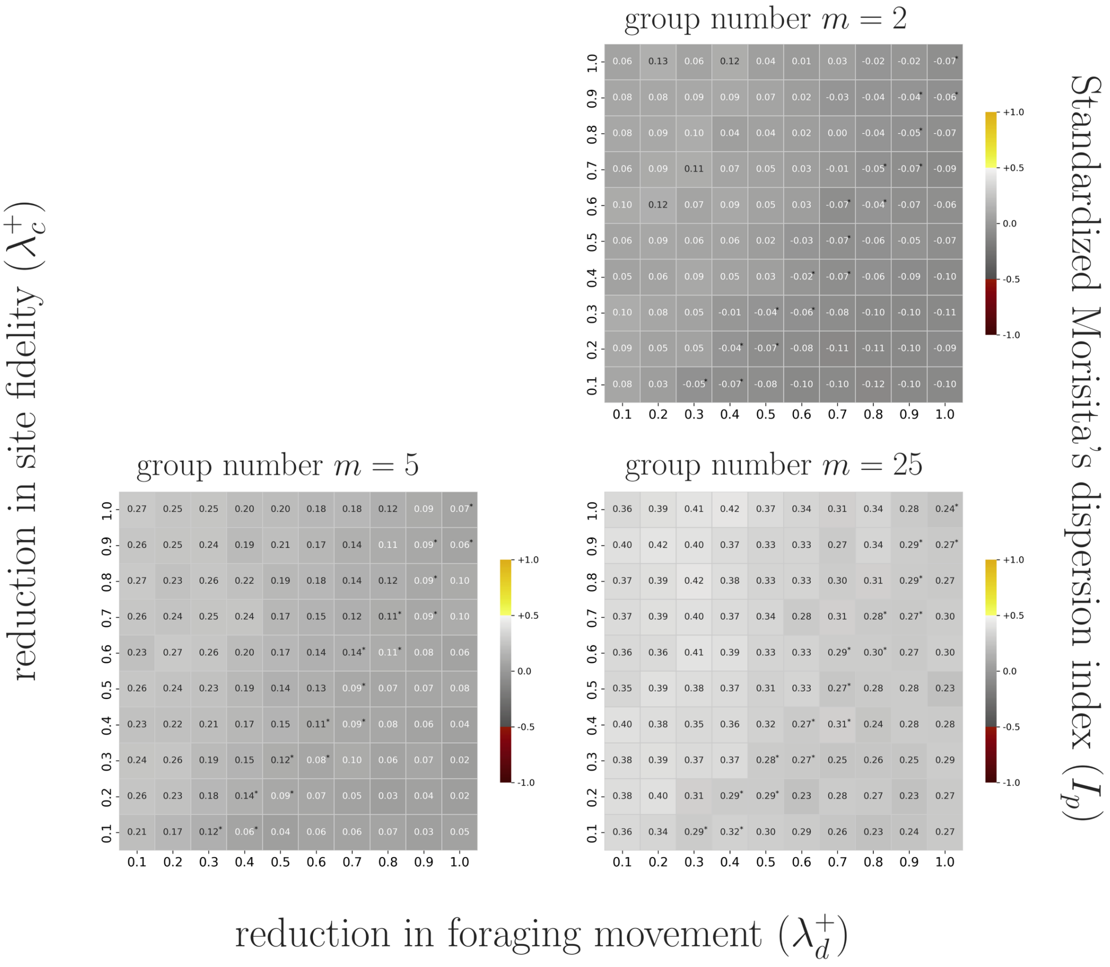


Supplementary Figure 2. As in Fig. S1, but corresponding to Fig. 4, with all parameters as in Fig. 4.

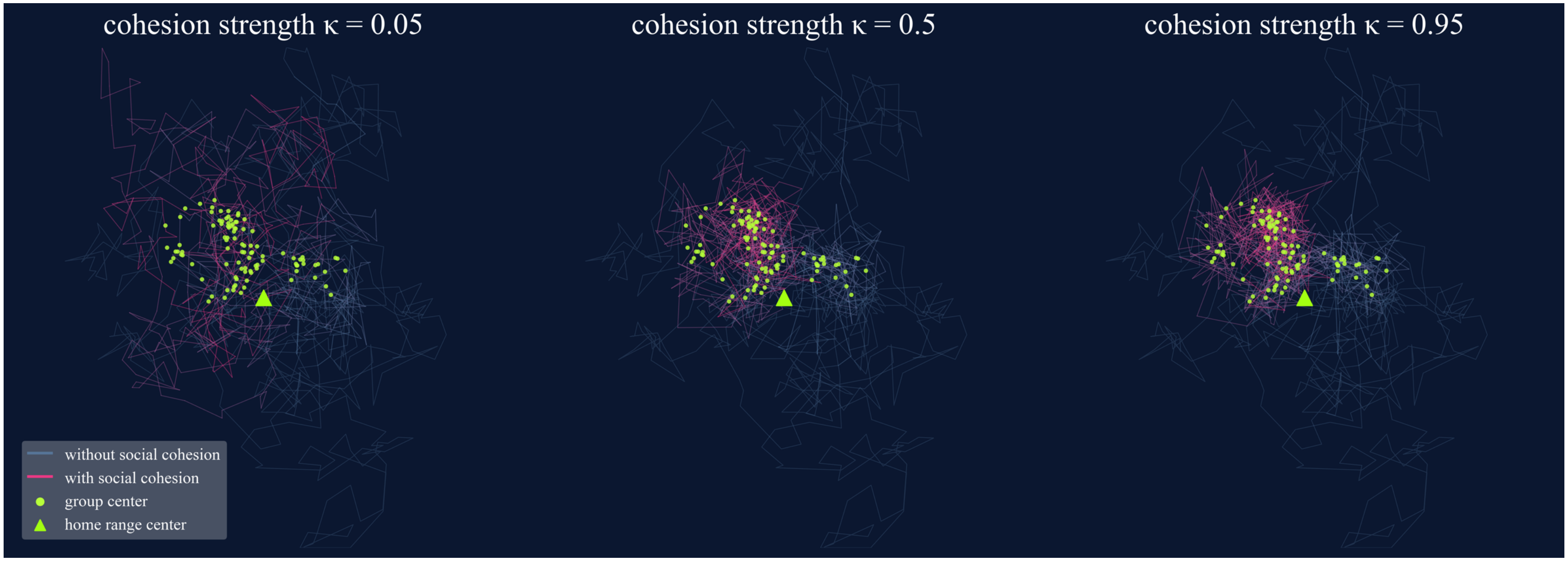


Supplementary Figure 3. Effect of social cohesion on simulated movement trajectories for five members of a single-group population. Gray lines show trajectories over the first 100 time steps, before cohesion is applied; purple lines show the subsequent 100 time steps under cohesion of strength $\kappa$. Green dots mark the group centroid at each time step (identical across $\kappa$, as all runs share a random seed); the green triangle marks the home-range center.

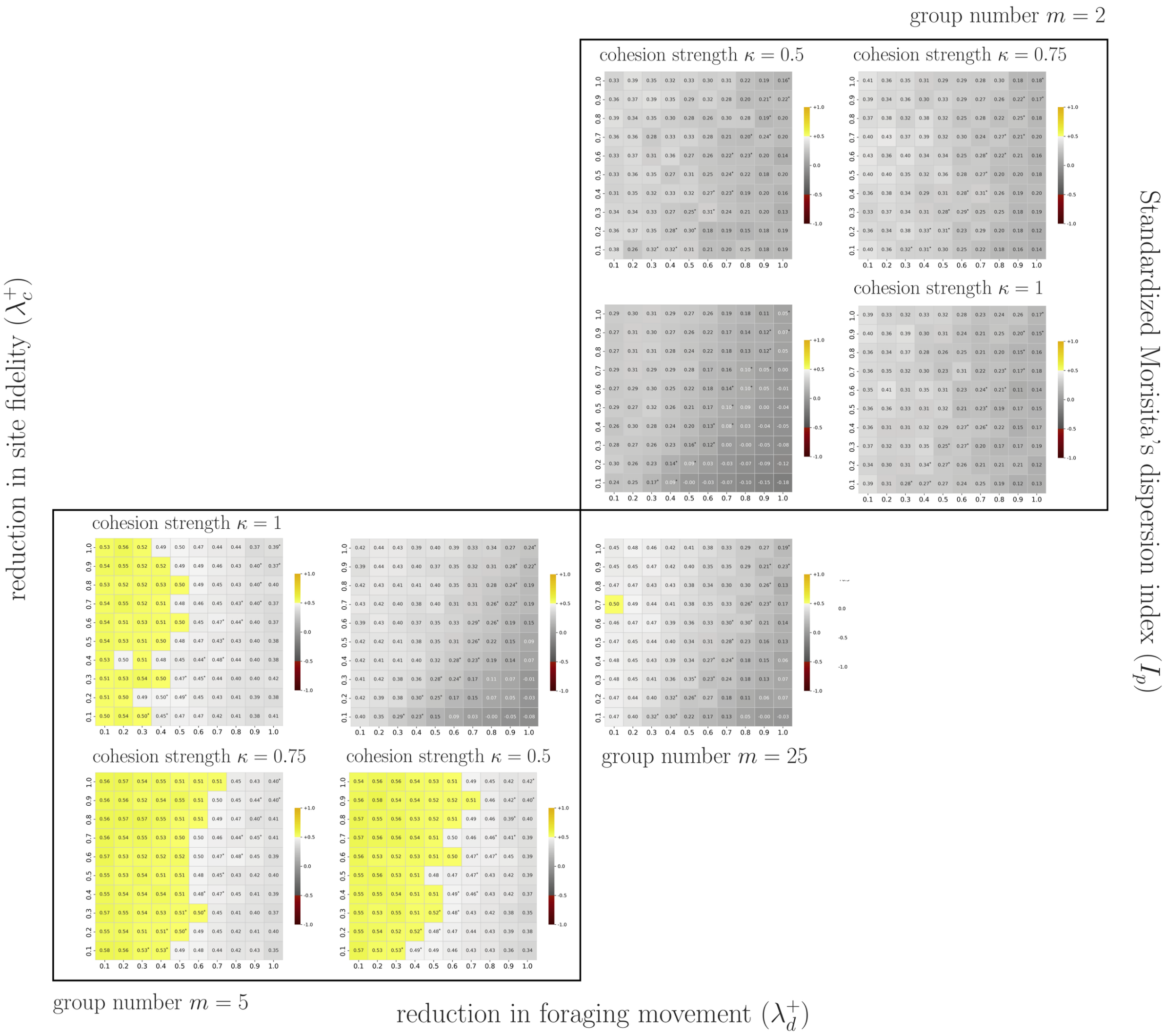


Supplementary Figure 4. As in Fig. S1, but corresponding to Fig. 6, with all parameters as in Fig. 6.

Supplementary Video 1. Animation of the simulation in Figure 3a, whose three panels show snapshots at selected time points.
